# Estimating age and annual tooth growth rates in narwhals (*Monodon monoceros*) using computed tomography

**DOI:** 10.64898/2026.08.27.747649

**Authors:** Evgeny A. Podolskiy, Genya Shimbo, Sung-Min Park

**Affiliations:** Arctic Research Center, Hokkaido University, Sapporo, Japan; Faculty of Veterinary Medicine, Hokkaido University, Sapporo, Japan; College of Dentistry, Dankook University, Cheonan, Republic of Korea

**Keywords:** narwhal, computed tomography, teeth, age, growth rate, Greenland

## Abstract

The spiraling tooth of the narwhal (*Monodon monoceros*) has long intrigued scientists, but the growth rates of its “tusk” are poorly known due to the difficulty to age narwhal. To learn more about the population dynamics and life history of this exploited species, previous studies have used sectioned teeth to age narwhals and calibrate other ageing methods based on aspartic acid racemization, radiocarbon or trace element analysis. However, few such studies have been conducted due to the method’s difficulty and the need to sacrifice the tooth. In this study, computed tomography (CT) was used on five teeth as a non-destructive means of estimating the age of narwhals, revealing different growth rates between erupted and embedded teeth during adolescence. Moreover, in combination with our reanalysis of all relevant literature, CT data clarify the annual growth rates of narwhal tusks, which peak before the age of 10 (*∼* 100 mm yr*^−^*^1^). This approach will be useful for investigating private and museum collections, and might also improve the use of trace element proxies in narwhal teeth and support sustainable harvest management.

## 1 Introduction

Growth layers in teeth are used to determine the age of terrestrial and marine mammals Hohn (2018). However, this methodology is challenging and controversial due to its destructive nature and disagreements between methods or readers, thereby limiting the biological insights gained from age data, particularly for the narwhal, *Monodon monoceros* (Hay, 1984; Lockyer et al., 2018; Read et al., 2018; Reiter et al., 2025b). The extraordinary helical tusk of the narwhal is actually a canine that is not located within its mouth, but usually erupts above the upper lip from the left side of the upper jaw (Nweeia et al., 2012; Turner, 1873). In the literature, this erupted tooth is referred to as a tusk (called *tuugaaq* by the Inuit). Accordingly, both terms are used interchangeably throughout this paper. The tusk is observed almost exclusively in males and can reach nearly 3 m in length. The right-side canine erupts rarely and remains embedded (or impacted), measuring *<*30 cm in length (Hay, 1984; Watt et al., 2020). Hereafter, we refer to the unerupted tooth as an embedded tooth, although some papers also refer to it as a tusk (Tyler et al., 2008). Both teeth have so-called growth layer groups (GLGs) that are expressed as light and dark growth increments in longitudinally sectioned teeth. A GLG refers to a group of layers that occur with cyclical, predictable repetition and can be used to estimate an animal’s age (Perrin and Myrick, 1980). It has been established that the GLG deposition rate in the teeth of the beluga (*Delphinapterus leucas*), which are the closest living relative to the narwhal, occurs in annual increments (Lockyer et al., 2018). In narwhal, the rate has also been assumed to be annual (Garde et al., 2012; Watt et al., 2020), which has recently been validated using the radiocarbon (^14^C) nuclear bomb pulse (Garde et al., 2024).

Embedded teeth (called *tugarrusirq* by the Inuit), have been investigated and shown to have GLGs in both calcified tissues of the teeth, dentine and cementum (Dietz et al., 2004; Hay, 1984; Teilmann and Dietz, 1993; Watt et al., 2020). New dentine layers are deposited on the internal surface of the tooth, whereas new cementum layers are deposited on the external surface. In narwhal, erupted tusks have GLGs in the dentine, but these are unclear in the cementum (Garde et al., 2012). GLGs in both teeth have been used to calibrate and compare the asparic acid racemization (AAR) rate, another method for estimating narwhal ages (Bada et al., 1983; Garde et al., 2012, 2007; Watt et al., 2020). The AAR technique uses some of the most stable proteins in the body, found in teeth and the nuclei of the eye lenses, and relies on the conversion of aspartic acid between its L- and D-forms in metabolically inactive tissues at a predictable rate, a process known as racemization (Garde et al., 2012). AAR analysis of embedded teeth and eyes enables the ageing of both males and females, with the latter usually lacking any erupted tusks. Paired embedded teeth, and a matching erupted tusk and embedded tooth yield consistent age estimates (Hay, 1984; Watt et al., 2020). However, the dentine of the embedded teeth is unusable for ageing narwhals older than 20 years due to closure of the root by cementum after reaching sexual maturity (Hay, 1984; Watt et al., 2020). After rapid elongation of the tooth and filling of the pulp cavity with four-five dentine layers in young narwhal, dental deposition decelerates, making the dentine GLGs narrow, compressed, and difficult to count (Hansen et al., 1990; Hay, 1984). As the enveloping cementum covers the entire root, deposition ceases (i.e., occlusion). Moreover, an additional problem is that a portion of the root or “knot” may remain in the skull during removal of the embedded tooth (Hansen et al., 1990; Teilmann and Dietz, 1993). Cementum did not appear to be useful for ageing narwhal (Hay, 1984). However, Dietz et al. (2004); Teilmann and Dietz (1993) investigated the dentine and cementum on the tip of the embedded teeth of narwhals and showed that the deposition of cementum GLGs also stopped after approximately 20 layers. Therefore, the embedded teeth allow us to determine only the minimum age of a narwhal.

In general, current methods of narwhal age estimation are either invasive (AAR and GLGs; Dietz et al. (2004); Garde et al. (2024, 2012, 2007); Watt et al. (2020)) or relative and imprecise (e.g., body whitening with age, or a GLG/length derived from Gompertz growth curve (Dietz et al., 2004; Garde et al., 2007; Lockyer et al., 2018)). Moreover, ageing methods based on teeth analysis require sectioning, segmentation, and sample preparation, followed by either challenging manual observations by an experienced tooth reader or sophisticated laboratory measurements such as radiocarbon measurements or laser ablation inductively coupled plasma-mass spectrometry (LA-ICP-MS) trace element analysis Dietz et al. (2004); Garde et al. (2024); Reiter (2026); Reiter et al. (2025a,b); Watt et al. (2020)). Furthermore, aligning the signals between consecutive tusk pieces with overlapping GLGs requires nontrivial procedures based on modeling (Reiter et al., 2025a,b). The corresponding destruction of a precious tusk, along with the time-, labor-, and resource-intensive analyses, makes narwhal age estimation difficult or even impossible when dealing with museum specimens, often due to a lack of funding or personnel. However, such age estimation is essential for understanding the basic biology and ecology of the narwhal, and its catch composition and stock demography, which is required for stock management and food security via regulations and quotas (Dietz et al., 2021, 2004). Therefore, this information is also of fundamental importance to Inuit hunters in Greenland and Canada, who harvest narwhals under quotas (Garde et al., 2012; Lockyer et al., 2018).

Dental research on other modern and fossil odontocetes using micro-computed tomography (*µ*-CT) suggests that three-dimensional studies of teeth and their internal morphology might be useful for non-destructive and non-invasive age determination for rare or valuable specimens (Loch et al., 2013). In the latter study on extant dolphins, grayscale variations in the dentinal layers were insufficient to clearly resolve GLGs. More recently, however, two harbor porpoises were successfully aged using *µ*-CT methods (Baier-Stegmaier et al., 2023). Since GLGs of narwhals are an order of magnitude thicker (hundreds of microns for dolphins versus 1.84 mm for narwhals; (Garde et al., 2012; Loch et al., 2013)), we hypothesize that they might be easier to resolve.

The purpose of this study was twofold. First, to show that CT is a promising method for aging tusked narwhals, as it does not require bisecting their expensive and difficult-to-obtain tusks; and, second, to estimate teeth annual growth rates, which remain unclear, as we show by a comparative analysis of all published results to date. Specifically, we present the following. A scan of a narwhal’s tusk takes *<*15 minutes to complete, which is followed by opening the raw data in open-source software and visually tracing the GLGs along the entire CT scan of the tusk on the monitor. This enables a reasonable estimation of a narwhal’s age within 1 h. In addition, mapping the identified GLGs along the length of the tusk, along with those in the matching embedded tooth, can for the first time provide biological insights into annual growth rates. Finally, the CT data presented here (Podolskiy et al., 2026) provide an open-source reference material for those in the scientific community who are investigating or teaching the unique dental and skull anatomy of the narwhal.

## 2 Materials and methods

### 2.1 CT scan of narwhal teeth

The target specimens of this study were two matching erupted (NW2203a) and embedded (NW2203b) teeth from a male narwhal (with an embedded tooth inside the skull), and one erupted (NW24) and two embedded (NW2313, NW2403) teeth from another three narwhals, all of which were harvested near Qaanaaq, northwest Greenland. Table 1 shows the specifics of all the specimens.

**Table 1.** Data for five teeth from four narwhals harvested in Inglefield Bredning, northwest Greenland, that were used in this study.

|  | Narwhal NW2203 |  | Samples from 3 different narwhals |  |  |
| --- | --- | --- | --- | --- | --- |
| ID no. | NW2203a | NW2203b | NW2313 | NW24 | NW2403 |
| Harvest date | 31/7/2022 | same | 21/8/2023 | 2024 | 9/8/2024 |
| Sex | M | same | M | M | M |
| Body length (cm) | 390 | same | 408 | unknown | 405 |
| Total tooth length (cm) | **155.5 (L)** | **20.0 - emb. (R)** | **21.4 - emb. (R)** | 133.5 (cut **124.5 (L)**) | **22.1 - emb. (R)** |
| External tusk length (cm) | 122.5 | same | 151 | 101.5 | 168 |
| Diam. of pulp cavity (mm) | 18 | 1.3 | 0.9 | 14.6 | 0.8 |
| Diam. of tooth (mm) | 42 | 18 | 19 | 39.2 | 18 |
| Tooth weight (g) | 1860 - tusk | unknown | 65 - emb. | 1450 - tusk | 62-emb. |
| Neonatal line present | yes | yes | yes | yes | yes |
| Tip of tooth present | yes | yes | yes | yes | yes |
| Base of tooth present | yes | yes | yes | no | yes |
| No. of resolved GLGs | 19 | 10 | 6 | 15 | 6 |

Note: NW2203a and NW2203b are the matching erupted tusk and embedded tooth of the same narwhal (NW2203). The analyzed tooth specimens are shown in bold. “L” and “R” indicate left and right. For the erupted tusks, the diameters of the tooth and cavity were measured approximately at the point of eruption from the maxilla. For the embedded teeth (or “emb.”), the diameter of the tooth was measured immediately before the knot at the root. The cavity diameter was measured at its location.

Animal sacrifice was not part of the study, as all narwhals had been previously harvested by licensed Inuit hunters for traditional subsistence purposes and not for research. Therefore, no approval from an ethics board or Institutional Animal Care and Use Committee was required for the bone scanning conducted in this study. The Government of Greenland permitted the export of the specimens to Japan (CITES Certificate Nos. 24GL2089848, 25GL2090136, and 24GL2089894).

The full skeleton of a male narwhal (NW2203) was purchased from Inuit hunters who harvested it in the center of Inglefield Bredning on 31 July 2022. The specimen was cleaned, degreased, and legally transferred to Hokkaido University, Sapporo, Japan, in 2025. Since March 2025, it has been stored at room temperature in the Hokkaido University Museum (HoUM; under ID HoUMVC-14735). On 12 February 2026 the narwhal head pieces, including the erupted tusk, skull, lower jaw (i.e., mandible), and the embedded tooth of another specimen (NW2313), were transported to the Faculty of Veterinary Medicine, Hokkaido University, Laboratory for Computed Tomography (CT). After CT imaging, the specimens were transported to the Arctic Research Center laboratory for anatomic examination, assembly, and exhibition at the HoUM, which was opened to the public on 1 April 2026. On 10 March, an erupted tusk NW24 and an embedded tooth NW2403 (harvested in summer 2024 near Qaanaaq) from other collections in Japan were also scanned to reconfirm the general findings of this study.

To create digital two-dimensional images and three-dimensional models of the narwhal skull (including the embedded tooth NW2203b) and other teeth, standard medical CT data were collected using an 80-slice multidetector CT scanner (Aquilion Prime; Canon Medical Systems, Tochigi, Japan). The maximum object size that can be scanned at once is approximately 500*×*500*×*1100 mm^3^. The scanner produced 0.5 mm-thick slices every 0.3 mm in each scan: 5374 slices for tusk NW2203a (the scan had to be split into two sections to fit samples 2482 and 2892), 2012 slices for the skull with embedded tooth NW2203b, 1722 for the mandible and separate embedded tooth NW2313, 5752 for the tusk NW24 (this scan also had to be split into two sections to fit samples 2871 and 2881), and 811 for the embedded tooth NW2403. Overlapping scanned sections of long erupted tusks were identified from length measurements and excluded to avoid double-counting of GLGs.

For the skull scans, the tube voltage and current were 120 kV and 500 mA, respectively. For the tusks and lower jaw scans, the settings were 120 kV and 350 mA, respectively. The rationale for these CT scanning parameters was that the specimens were *ex vivo* osseous structures, where radiation exposure was not a factor, and parameters could be selected to prioritize image quality and spatial resolution. A tube voltage of 120 kV was used to ensure adequate X-ray penetration of the large, highly attenuating bony structures. The maximum available tube current was used for each scan condition to reduce image noise, as increasing the X-ray dose decreases quantum noise in CT images. In addition, a small focal spot was used whenever technically feasible to maximize geometric sharpness and improve spatial resolution. For the skull scans, the large specimen size required a wide field of view, precluding the use of the small focal spot. Therefore, a tube current of 500 mA was used. For the tusk and lower jaw scans, the smaller specimen size allowed the use of the small focal spot, for which the maximum available tube current at 120 kV was 350 mA.

All scans were reconstructed on a 512*×*512 pixel matrix. For the skull, we used a field of view (FOV) of 475 mm, and a focal spot size of 1.6/1.4 were used. For the erupted tusks (NW2203 and NW24) and one embedded tooth (NW2403), the scans were performed with a FOV of 150 mm, and a focal spot size of 0.9/0.8. For the lower jaw and separate embedded tooth (NW2313), a FOV of 306 mm and a focal spot size of 0.9/0.8 were used. The data were stored as Digital Imaging and Communication in Medicine (DICOM) files (Podolskiy et al., 2026). The physical units used in the files are millimeters for linear dimensions and Hounsfield units (HU) for CT attenuation values that relate to the radiodensity.

Previously, both sides of the longitudinal section of the erupted tusk have been used for GLGs readings (Garde et al., 2012). While tracking the longitudinal propagation of each GLG by eye is challenging (Reiter, 2026), it is simpler on CT images using image tools. Instead of counting the GLGs as bands on longitudinally bisected erupted or embedded teeth (as commonly done (Garde et al., 2012; Hay, 1984; Watt et al., 2020)), their appearance was visually tracked in a CT cross-section of a tooth. Such cross-section reminds tree rings, and by moving between the tip and root along the axis of the tooth, the reader can track cones of GLGs as newly emerging rings, whilst reading the corresponding distance from the tip on the screen. This approach is less subjective and differs fundamentally from the destructive GLG counting method because it avoids the main sources of error associated with having one or more readers examine the same irreversibly bisected, polished and chemically prepared specimen under varying light conditions (Hohn, 2018; Watt et al., 2020). Therefore, it remains to be determined how the established criteria for ageing monodontids (Lockyer et al., 2018; Read et al., 2018), recently shown to be imperfect (Reiter et al., 2025b), can be adapted to the CT approach; addressing this issue, however, is beyond the scope of the present study.

Furthermore, to our knowledge, no previous study that relied on visual counting of GLGs in narwhal teeth included multiple age readers and provided error estimates like in other marine mammals (Read et al., 2018). Instead, previous studies relied on a single, experienced tooth reader trained in reading beluga teeth or embedded narwhal teeth (not tusks), because less experienced readers are known to be negatively biased (Garde et al., 2012; Hay, 1984; Watt et al., 2020). However, no experienced readers are available for reading the CT scans of a narwhal, especially in cross-sectional view. Here, as in previous studies, a single reader (EAP) was chosen to read the data for consistency. While there is no universal optimal number of readings per tooth, at least 3 is recommended (Read et al., 2018). Therefore, the reader viewed all the teeth during two sessions in March 2026 and one session in August 2026. During the first session, the reader missed all compressed GLGs after finding six GLGs in the embedded tooth NW02203b due to its bending near the root. Subsequently, three consistent and unambiguous age readings were achieved. Co-authors could not find further GLGs, however, they knew the count from personal communication with EAP and could not be considered independent. Other readers were sought at the international symposium on narwhal tusks on 8-10 June 2026 in Copenhagen, but no-one had time to contribute independent readings.

The files were processed in Weasis v. 4.6.6 and 3D Slicer v. 5.10.0 for the two- and three-dimentional reconstructions, respectively, on a MacBook Pro laptop with 32 GB of RAM. The Segment Editor in 3D Slicer was used to isolate and extract structures for visualization. We relied on the fact that teeth have a higher density than the skull and other tissues, allowing a so-called threshold-based segmentation, which allows “removing” teeth from the skull for visualization.

The adequacy of the CT approach for age estimation can be independently assessed by comparing the resulting age estimates with expected GLG deposition rate. To validate the age estimates derived from CT and compare them with the most widely recognized ageing methods – counting GLGs by naked eye after bisection and using AAR – we compared our results with published age estimates, as outlined in the next section.

### 2.2 Comparative analysis of growth rates

For erupted and embedded teeth, length data are limited to a few studies (Dietz et al., 2021; Garde et al., 2012, 2007; Hay, 1984; Reiter et al., 2025b; Watt et al., 2020), all of which only allow indirect estimates of the average growth rate by dividing the reported length of the tooth by the estimated age. For the erupted tusks, growth rates derived in this way can be underestimated by using the external length instead of the rarely available total length (i.e., obtained after extraction from the cranium, which can be up to 39 cm longer; e.g., sample 956 in Garde et al. (2012)). The lengthening rate can also be overestimated by ignoring the fact that narwhals are usually born with 9.5 cm of unerupted tooth (Hay, 1984) (which is also crucial for individuals with short embedded teeth). For these reasons, we first summarize the available records and adjust these to enable a consistent comparison, as explained below.

#### 2.2.1 Erupted teeth (tusk)

Hay (1984) analyzed data from narwhals harvested in the eastern Canadian Arctic and compared the external length of 52 unbroken male left tusks and 1 unbroken female tusk with the number of growth layers in dentine and the mandible. Their study suggested that the tusk erupts at *∼*3.5 growth layers and attains a maximum external length of *∼*215 cm at 50 GLGs.

For 26 male narwhals with measured external tusk lengths (harvested in Greenland), Garde et al. (2007) estimated their age using the AAR method and provided a fit using the von Bertalanffy growth model, which showed young narwhal had a faster growth rate than older individuals, as follows:

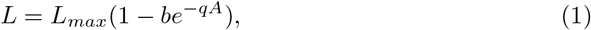

where *L_max_*is the asymptotic standard length of 178 cm reached at approximately 53 years of age, *A* is the age estimate, and *b* and *q* are model constants (1.197 and 0.0921, respectively). The derivative of Eq. 1 gives the growth rate (cm yr*^−^*^1^) as follows:

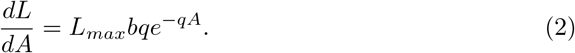

This relationship yields an instantaneous or age-specific growth rate. As such, it is not straightforward to compare the result with the averaged growth rate obtained from the year of death (*Ĝ* = *T_L_/A*), which might obscure potential growth nonlinearities. Therefore, instead of plotting as it is, the tusk length reached at certain age *L*(*A*) was divided by the corresponding age *A*. There are two ways of doing this, both of which extend beyond the intended range of the original fit (Garde et al., 2007) and involve dividing Eq. 1 by the corresponding age or back-integration of the derivative Eq. 2:

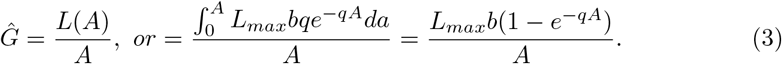

The first approach yields negative growth by an age of 2 years, and the second method overestimates growth by the age of 2 years due to a steep gradient from 0 to 2 years. Based on this, the first two years were ignored (i.e., negligible growth was assumed) and the first, simpler approach was used for further comparisons (referred to herein as the modified von Bertalanffy model).

Furthermore, the external tusk length and age data produced by Garde et al. (2007) (their appendix I) were subsequently converted to average growth rates and plotted on a logarithmic scale by Nweeia et al. (2008). This previous study did not examine the implicit growth rate relationship in Garde et al. (2007) as done in the present study (see above), but it highlighted the decrease in growth rate with age by fitting a regression, which yielded R^2^ = 0.7517, but missed the smallest measurement (on whale No. 557). Our regression of the original data (Garde et al., 2007) yielded the same R^2^ value only after multiplying the missing measurement by 10. For the correct data, our R^2^ is about half of this value (0.3757); however, this relationship (in mm yr*^−^*^1^) was used in a non-logarithmic form for convenience and to enable comparisons:

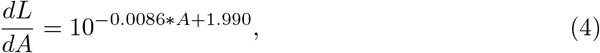

Garde et al. (2012) studied Greenlandic narwhals and obtained their life-averaged growth rates by dividing the total length of each of 11 erupted tusks (at least one from a female) by the matching number of visually identified GLGs (mean = 63.3 mm yr*^−^*^1^; ranging = 35.4–91.1 mm yr*^−^*^1^). The lowest rates corresponded to the oldest tusks (60–70 GLGs), suggesting that growth rates likely ceases with age. According to our reanalysis, using the external tusk length leads to a lower mean of 47.1 mm yr*^−^*^1^. More recently, Dietz et al. (2021) counted dentine GLGs in 10 tusks (seven unbroken) from Northwest Greenland male narwhals to explore their lifelong feeding ecology. We refer to their Table 1 and, for three broken tusks, use the reported length and the total estimated age (by adding the mean of missing years in the tusk tip). The unknown length of the broken tusk tip might lead to an underestimation of the growth rates by 0.3–0.6 cm per year. Finally, Reiter et al. (2025b) presented age estimates for 16 narwhals from Greenland with known total tusk lengths including 4 females (some of these narwhals were previously analyzed by Garde et al. (2024)).

In calculating the growth rate from these previous data, the total tusk length was reconstructed, if unknown, by adding the hidden part (*L_h_*) using a regression fit. This regression was produced using data from Hay (1984) and Garde et al. (2012) with *R*^2^ = 0.8078 and had the following form:

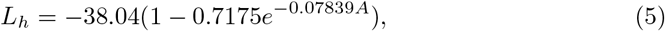

The presumed tooth length at birth of 9.5 cm (Hay, 1984) was then subtracted from the total tusk length and finally divided by the estimated age.

#### 2.2.2 Embedded teeth

Watt et al. (2020) and Garde et al. (2024) published matching embedded teeth lengths and age estimates for 20 and 8 narwhals (of which 18 and 6 were females), respectively. After subtracting the presumed tooth length at birth (9.5 cm (Hay, 1984)) from the reported length of each specimen and dividing it by the mean number of counted GLGs, the life-averaged growth rates were obtained.

## 3 Results

### 3.1 Qualitative assessment

The data for narwhal sample NW2203 revealed fine details of the skull, including an embedded tooth (20 cm long), the socket of the main tusk with a depth of 33 cm, an embedded vestigial right tooth (4.5 cm long), and an asymmetrical skull structure (Fig. 1). The socket of the left vestigial tooth was also observed, but the tooth was not scanned as it fell out during sample preparation and cleaning. CT images of the erupted and embedded teeth show the pulp cavity, cementum, and distinct layers in dentine corresponding to growth rings in a cross-sectional view. These dentinal GLGs appear as elongated cones that can be truncated at the pulp cavity. Note that while the orientation of longitudinal CT section of the erupted tusk strongly affects the apparent position of the GLGs, particularly the pulp cavity, CT-data can be viewed and inspected from any angle on a screen. All five scanned teeth, including the embedded tooth inside the skull (NW2203b), show GLGs. The diameter of the vestigial tooth (4.5 mm) was too small to resolve any layers in dentine, whereas an earlier study reported none (Nweeia et al., 2012). Individual cementum GLGs could not be resolved at the current scan resolution. All embedded teeth had knots (e.g., label “2” in Fig. 1) that are known to appear at their root in male narwhals after seven dentinal layers (Hay, 1984). CT imaging also allowed visualization of the spiraling tusk morphology, which is observed as a spiral pattern of grooves and ridges not only on the outer tusk surface, but also along the interface between the dentine and pulp cavity, and on the surface of the socket containing the tusk (Fig. 1). CT scans of both tusks showed surface radial cracks that are up to 5 mm deep and propagate along the spiral, which were caused by drying of the specimen (Fig. 1).

**Fig 1.**
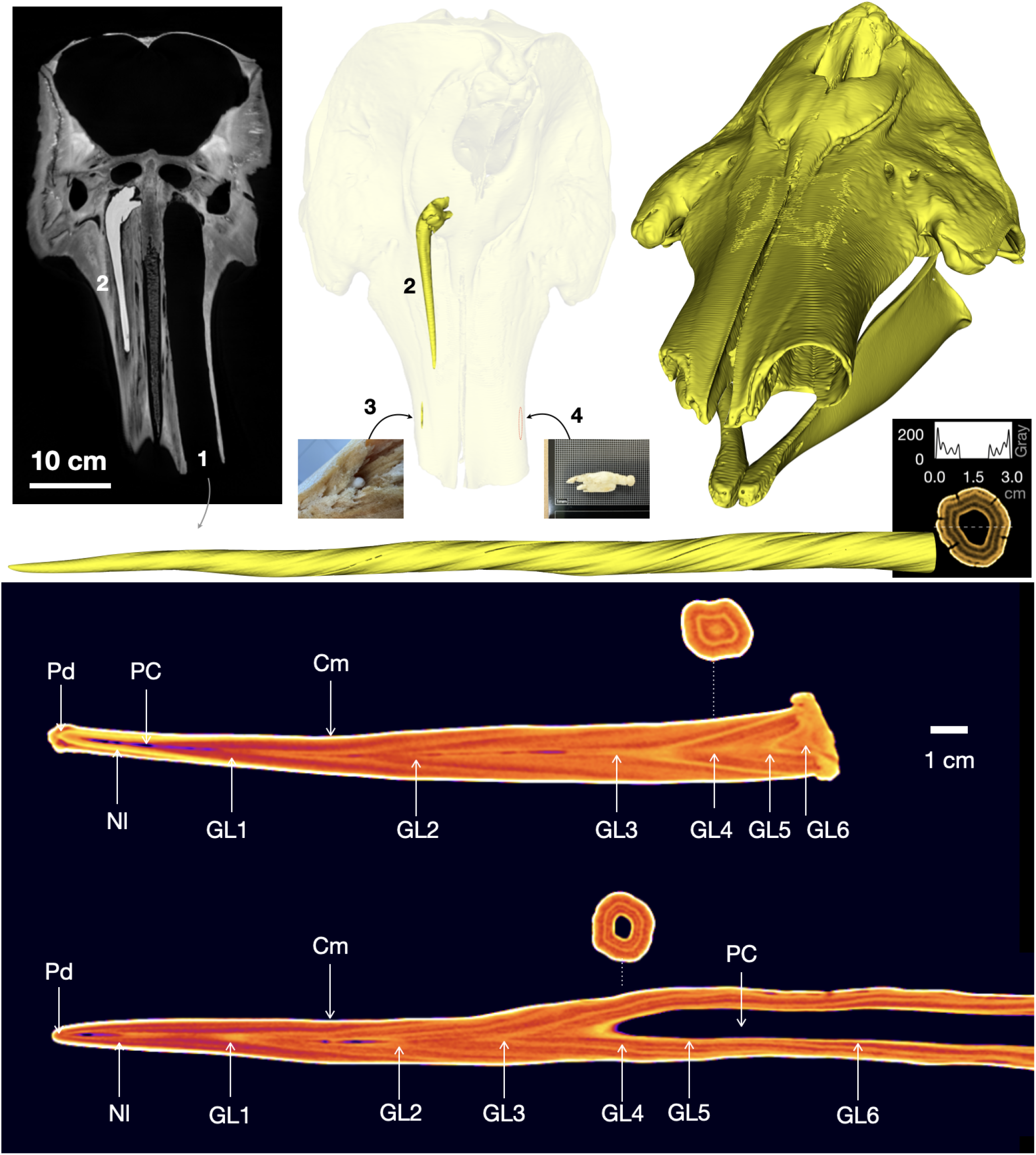
Dental anatomy of a narwhal with CT sections and 3D reconstructions. Coronal view of a narwhal skull shows (1) the socket of the left erupted tooth, (2) an occluded embedded tooth (NW2203b), and (3–4) a pair of vestigial teeth. The CT cross-section highlights the layering using grayscale values for a transect across the middle of tusk NW2203a. Longitudinal median CT sections show an embedded right tooth (NW2403) and apical tip (distal part) of the left erupted tooth (NW2203a), with the abbreviations following (Hay, 1984) and (Garde et al., 2012): Cm = cementum, NI = neonatal line, GL = growth layer band, Pr = prenatal dentine, and PC = pulp cavity.

### 3.2 Quantitative assessment

Examples of the location of GLGs along the erupted and embedded teeth are shown in Fig. 2. Six to 10 dentinal layers with declining thickness in all three embedded teeth were resolved. All teeth showed the greatest increase in length after the second layer (evident in Fig. 1 as the largest increment between GLGs 2 and 3), which was followed by a decrease in growth rate. In the erupted tusks, 15 and 19 layers were resolved, which in contrast to the embedded teeth, showed increasing thickness after an age of 4–6 years. It was not possible to resolve nucleation of new GLG cones within the socket section of both erupted tusks (*∼*30 cm from the root). After disappearance of older cones, the layer of the latest cone compresses as the wall thickness decreases towards zero in the intact tusk NW2203a, and corresponds to a very thin and fragile fringe. The length increment per growth layer (i.e., the annual growth rate) of the erupted and embedded teeth were similarly low during the early postnatal years and diverged thereafter. The tusk (NW2203a) had more GLGs than the matching embedded tooth (NW2203b), apparently due to insufficient CT resolution in the thinnest, most compressed layers near the knot, as detailed in the Discussion. The erupted tusks NW2203a and NW24 had five and four twists, respectively (counterclockwise when viewed from the tip), corresponding to approximately one twist every 3.8 years. However, none of the embedded teeth had a complete twist.

**Fig 2.**
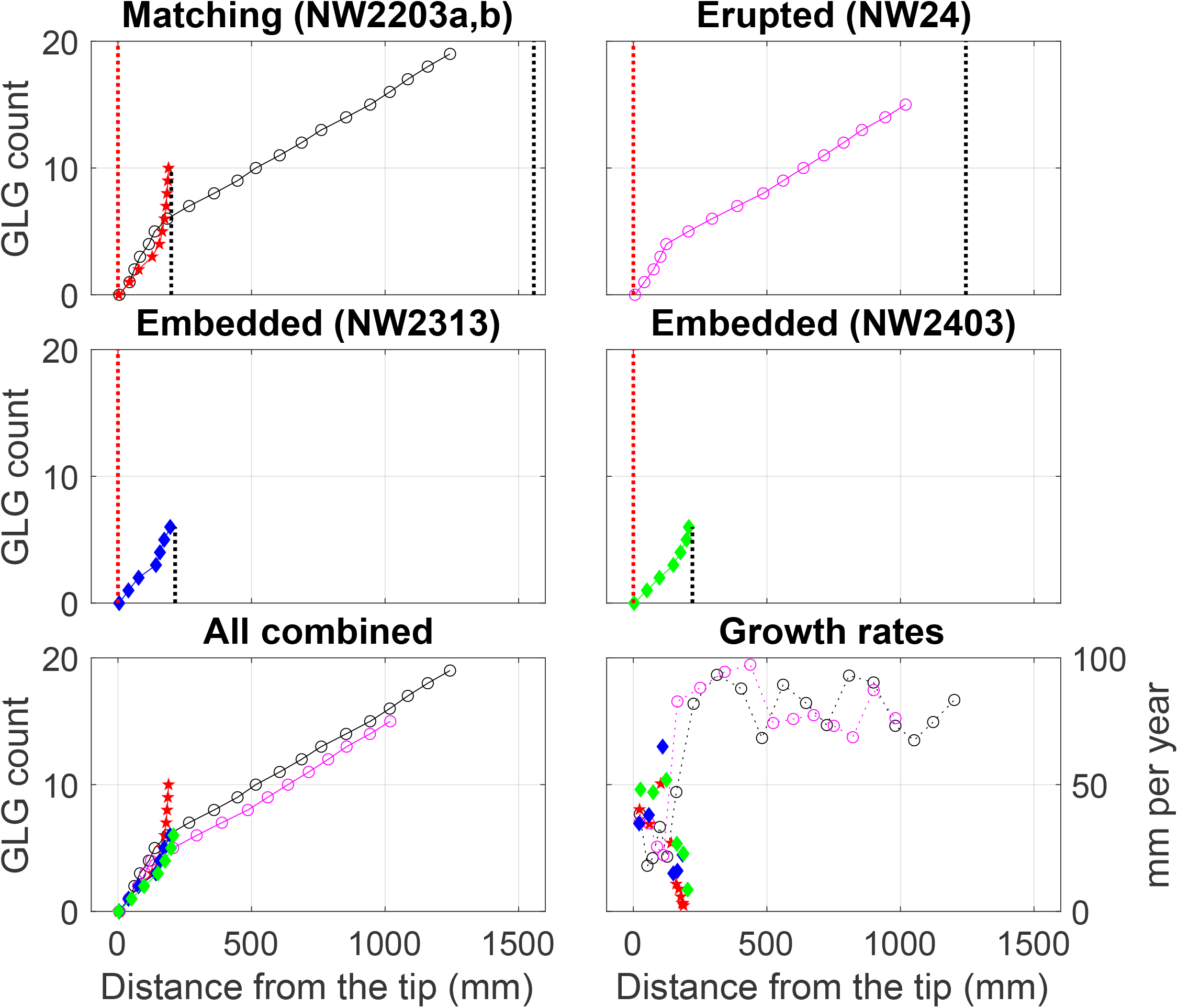
The number and the nucleation location of GLGs along the teeth axes obtained from CT imaging. Red and black dashed lines show the tip and the root of each tooth, respectively. Only NW2203a and NW2203b are matching teeth (black and red markers); other teeth belong to different narwhals (magenta, blue, and green markers). Lower subplots show all specimens, with their corresponding growth rates.

Combining the measurements from the previous studies (Dietz et al., 2021; Garde et al., 2012, 2007; Hay, 1984; Reiter et al., 2025b) and this study for all erupted tusks (*n* = 53 + 26 + 11 + 16 + 10 + 2 = 118) results in the most comprehensive overview of tusk lengthening with age to date (Fig. 3).

**Fig 3.**
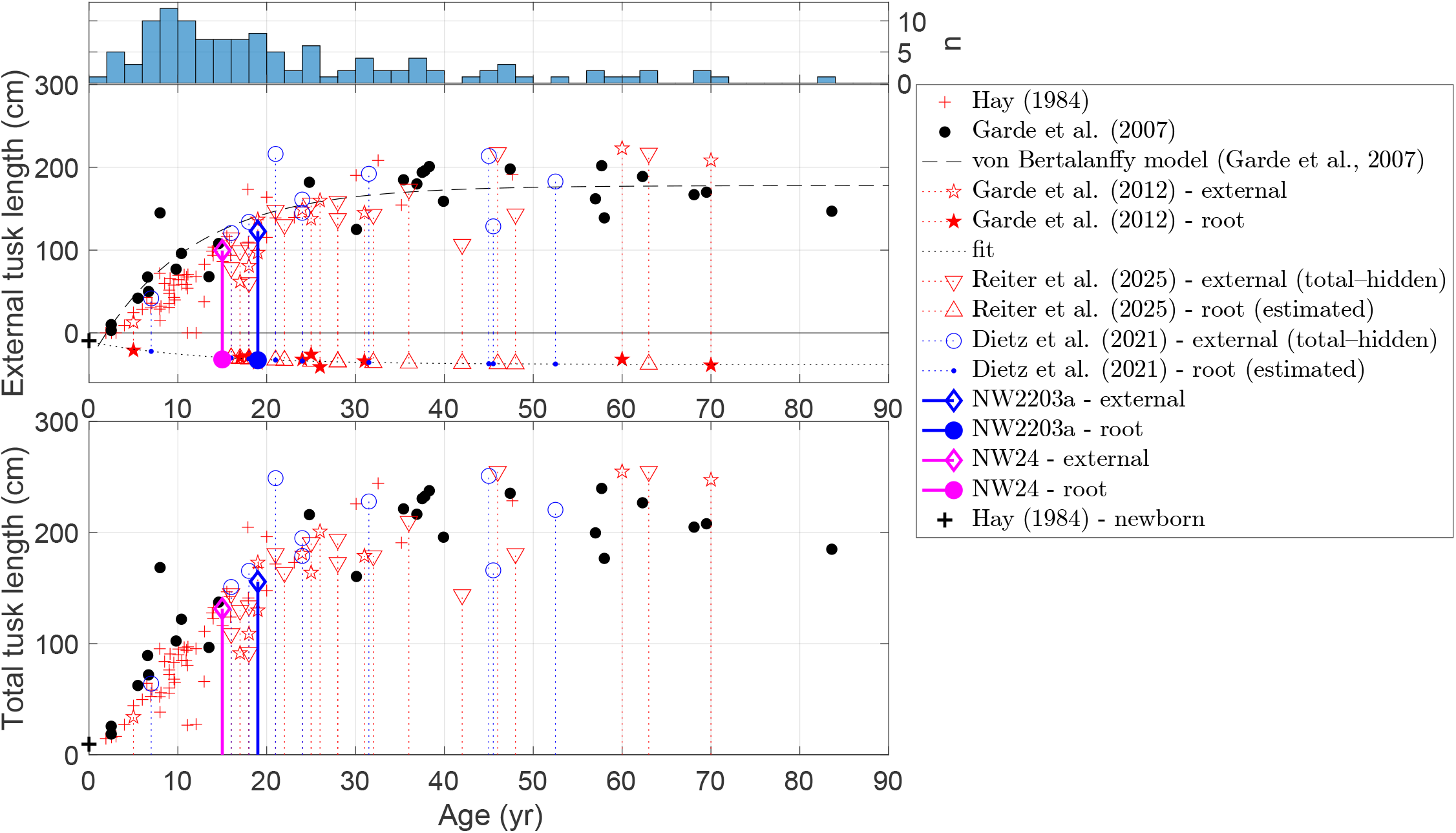
The CT-scanned tusks NW2203a (in blue) and NW24 (in magenta) are compared with literature and model data on tusks with estimated age and reported length (Dietz et al., 2021; Garde et al., 2012, 2007; Hay, 1984; Reiter et al., 2025b). The histogram shows the estimated age distribution. The dashed line shows a von Bertalanffy model (Eq. 1) which was produced for the external tusk length as a function of age, based on Garde et al. (2007). Negative values of the external tusk length correspond to the known or estimated length of the hidden section of the tooth (i.e., the distance to the rostrum). Filled circles and red stars show the measured length of the unseen tusk section (this study and Garde et al. (2012); the only exceptions are their samples 4076 and 888, with unknown external lengths). The dotted line is a fit (Eq. 5) to the data from Garde et al. (2012); Hay (1984), which was used to reconstruct the total length when it was unknown. For data with only a known total length (Dietz et al., 2021; Garde et al., 2007; Hay, 1984; Reiter et al., 2025b), open symbols show the offset made by assuming that the hidden length follows the regression (Eq. 5). A plus symbol + at the zero age corresponds to the typical 9.5 cm length of an embedded tooth at birth (Hay, 1984), which is shown for reference.

Based on the CT-estimated ages of the tusks (NW2203a and NW24), their external and total lengths fall within the expected range for animals of the same age (Fig. 3). The corresponding growth rates are shown against the previously published data and models in Fig. 4. It is noteworthy that the von Bertalanffy model is a modification of the fit (Eq. 3) for the external tusk length as a function of age, as reported by Garde et al. (2007), whereas the regression of Eq. 4 was undertaken on the same data from Garde et al. (2007) by Nweeia et al. (2008). For consistency, the histograms show only one value per tooth, obtained by dividing each tooth length by the age estimate, yielding life-averaged values. The plotted data yield a mean growth rate of 66*±*25 mm yr*^−^*^1^ (one standard deviation) and a median of 69 mm yr*^−^*^1^. This corresponds predominantly to the young narwhals in the sample set (median age of 17; Fig. 3). The estimated annual growth rates derived from the previously published data agree overall with the directly measured annual growth rates of the present study. The rates are low during the first postnatal years, but accelerate abruptly in teenage narwhals.

**Fig 4.**
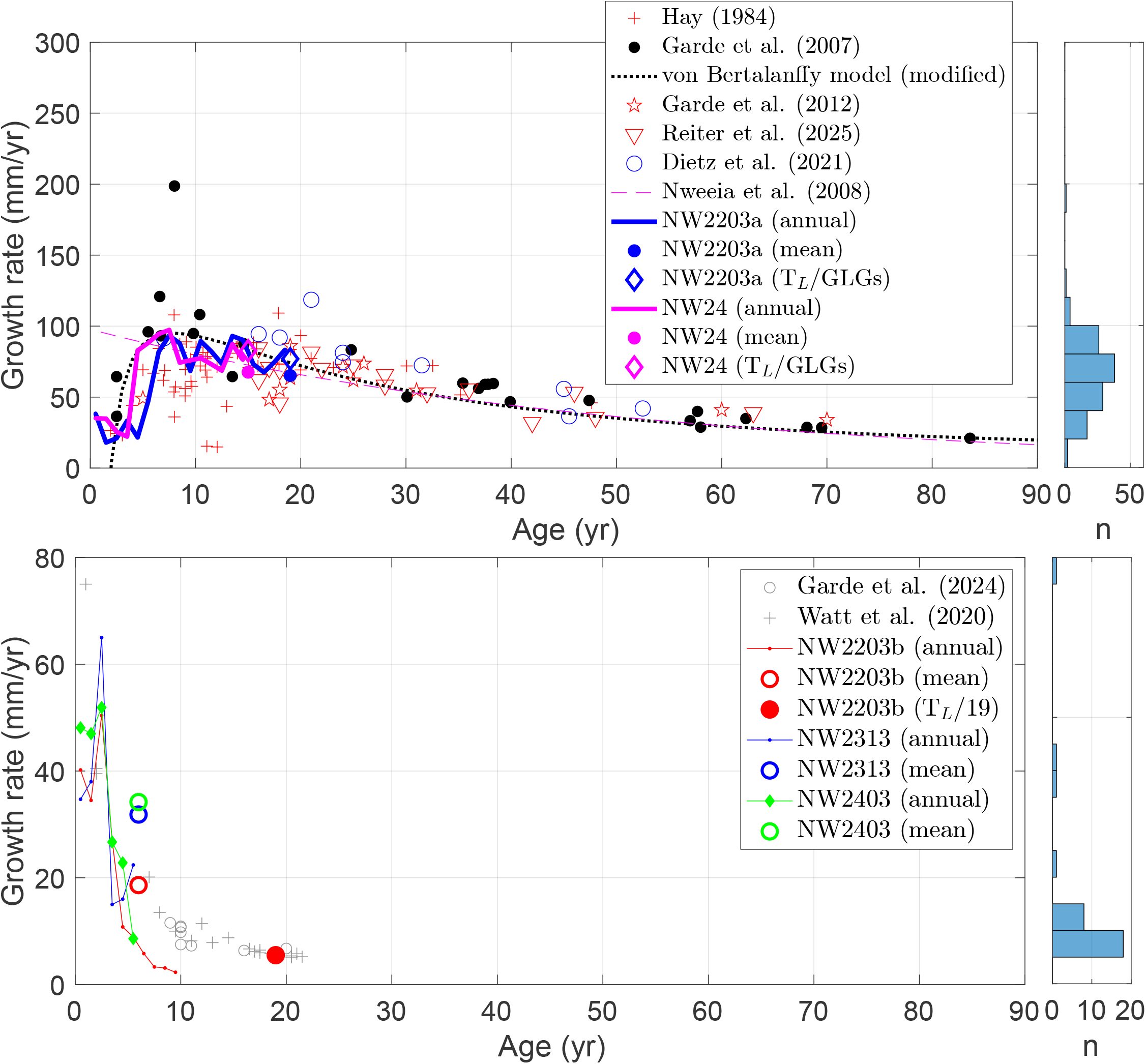
CT-derived annual growth rates of the erupted tusks NW2203a (in blue) and NW24 (in magenta) as compared with re-analyzed literature and model data (Dietz et al., 2021; Garde et al., 2012, 2007; Nweeia et al., 2008; Reiter et al., 2025b). The lower plot shows CT-derived annual growth rates for embedded teeth NW2203b, NW2313, and NW2403 (in blue, red, and green, respectively) as compared with data from the literature (Garde et al., 2024; Watt et al., 2020). See the text for details.

The life-averaged growth rates of embedded teeth derived from the literature (Fig. 4) appear to decrease exponentially over time. This is a manifestation of the decrease in the growth rate and cessation of GLG deposition in maturing narwhal (Hay, 1984), which implies that simple division produces somewhat misleading minimal growth rates, which must be higher in young animals, but can be very small after the first several years of life. In our study, the embedded teeth (NW2203b, NW2313, and NW2403) stopped lengthening after an age of 6. All the subsequent compressed layers, which were found in NW2203b but might have been unresolved due to their thinness in the other two samples (NW2313 and NW2403), no longer contribute to tooth lengthening. For example, dividing the length of embedded tooth NW2203b by the estimated age of the erupted tusk (NW2203a; age = 19 years) yields a comparable value to the data from other studies, which is lower than the mean of the CT-derived annual growth rate before an age of 6 years (red circles in Fig. 4).

## 4 Discussion

Previous studies considered the use of *µ*-CT imaging for non-invasive and non-destructive dental research of cetaceans, but were limited to two porpoises or suggested the need for refinements to enable aging research (Baier-Stegmaier et al., 2023; Loch et al., 2013). CT-based three-dimensional visualization of the dental anatomy of the narwhal was previously undertaken on one adult male, one adult female, and one female fetus, but was not focused on aging (Nweeia et al., 2012; Tyler et al., 2008). The visualization of that previous study, supported by examination of other skulls, showed (1) the presence of two vestigial teeth associated with evolutionary obsolescence, (2) that the pair of tusks and pair of vestigial teeth switch sides during development, and (3) that the tusks are canine teeth developed from the maxillary bone plate, as previously known (Turner, 1873). In contrast to the earlier studies, the novelty of the present study is the aging and growth-rate estimation, which is not based on the synthesis of data from many animals, but on individual narwhal.

To date, the age-specific growth rates of narwhal tusks remain poorly documented and have not been derived from comprehensive comparative analysis of literature or from previous von Bertalanffy model Garde et al. (2007). Therefore, the growth rates have been modeled in recent studies using stochastic processes without biological constraints (Reiter, 2026; Reiter et al., 2025b). Several other variants of the growth process have been proposed recently (gamma, Ornstein–Uhlenbeck, and piecewise linear), but have not been validated (Reiter, 2026).

Our results and comparative analysis show that the tusk annual growth rates are low during the first postnatal years, and accelerate abruptly in maturing narwhals. Therefore, this shows that the fit of Nweeia et al. (2008) overestimates the early-life growth rates, while the reanalysis of previous data and the von Bertalanffy model adapted from Garde et al. (2007) appear to yield similar results to those revealed by the CT data. The model fit intersects with the *x*-axis at 2 years, which is presumably the age of weaning (Reiter, 2026), before which an erupted tusk would be a disadvantage during the suckling period. As the CT results do not include narwhals older than 20 years, and based on other published data (Fig. 4), the rates are expected to decrease thereafter.

Hay (1984) provided the first analysis of embedded tooth growth and showed that tusks grow faster in young narwhals than in sexually mature individuals. They reported an average length of 21.67 mm for the embedded teeth of males and suggested that during early postnatal life, the tooth attains its maximum length after the deposition of six dentinal layers in males (four to five in females). Subsequently, the layer thickness decreases steadily, which explains the results of the present study (Figs. 2 and 4). More recently, Watt et al. (2020) showed that the dentine layer that closes the pulp cavity in the embedded tooth is deposited between 5 and 15 years of age. Therefore, the CT scan confirms that most of the dentinal GLG thickness is accreted in embedded teeth by an age of 6 years.

In general, the CT data show that the erupted and embedded teeth grow at similar rates until the age of around 4–6 (Figs. 2 and 4). Subsequently, the growth rates diverge. The erupted tusk accelerates (presumably until the age of 20 years) and twists, while the deposition rate of embedded tooth dentine decelerates, making GLGs unrecognizable in the CT images, and then ceases (Fig. 4). This remarkable deceleration occasionally fails, leading to anomalies such as double-tusked narwhals (0.9 *−* 1.03% of individuals in Greenland and Canada, respectively; Hay (1984); Louis et al. (2025)). Interestingly, a previous study on age determination using the ear plug in fin whales described an abrupt change in growth layer characteristics that coincided with the transition from sexually immature to sexually mature whale (Lockyer, 1984). We suggest that our data reveal a similar transition phase in narwhal teeth (Fig. 2).

Based on (i) studies that estimated age using the AAR and GLGs methods and (ii) our current understanding that 1 GLG/year is laid down (Garde et al., 2024), sexual maturity in male narwhals is attained between 9 and 17 years (Garde et al., 2007; Hay, 1984). Despite the uncertainty, it is likely that the tusk growth rates are highest around the period of sexual maturation, which is plausible when considering the likely importance of tusk length for breeding success (Graham et al., 2020).

Some variability between tooth readers is possible, and there are uncertainties associated with visual tracking and modeling outputs (Garde et al., 2012; Reiter et al., 2025b). However, instead of laboriously preparing sections to identify GLGs or undertake trace elements analysis, the CT method can be used to non-destructively age unique teeth or skull specimens. Importantly, since no cutting and chemical processing is involved, CT analysis of teeth is unaffected by issues like the location of the longitudinal cut (i.e., uneven sectioning), or incorrect staining and decalcification that can partly obscure the visibility of the layers in any tooth (Garde et al., 2007; Hay, 1984; Hohn, 2018; Watt et al., 2020). Moreover, open-source CT data will enhance the reproducibility of narwhal age studies. Such digital data can be readily re-examined (Podolskiy et al., 2026), whereas actual narwhal specimens are unlikely to be available to non-members of the laboratory or group undertaking the original assessment, due to the difficulty of export and import under the Convention on the International Trade in Endangered Species of Wild Fauna and Flora (CITES; also called the Washington Convention) or the termination of the original project. The accumulation of a larger number of CT scans will allow systematic uncertainty assessments to be performed to compare results from tooth readers and models. Also, the time-warping model of Reiter et al. (2025b) currently uses trace elements data as a proxy for annual cycles and age determination; this model is likely adaptable to grayscale value-distance profiles (Fig. 1; (Loch et al., 2013)).

Although female narwhals remain more difficult to investigate than males, *µ*-CT scans with significantly higher resolution and lower slice thickness (e.g., a 1 *µ*m pixel size (Baier-Stegmaier et al., 2023; Loch et al., 2013)) could be potentially useful for robust GLG counting in dentin or cementum of embedded teeth. Because *µ*-CT analysis of embedded teeth would not allow the ageing of narwhals older than *ca.* 20 years,

*µ*-CT investigation of the understudied mandibular periosteal layers could provide reliable age results (Hay, 1984; Teilmann and Dietz, 1993).

The regularity of the twist-to-age ratio raises the question as to whether a helical twist count could serve as an approximate supplementary age proxy, which warrants a systematic investigation of a larger sample set. The cracks running along the spiral were confined to the outer surface and did not intersect the central, dentinal GLG bands.

However, desiccation-related changes in HU (Hounsfield units) values near the tusk surface might be possible. As such, it is unclear how layering visibility might differ between recently harvested and old tusks (Loch et al., 2013).

Finally, in older individuals, early GLGs can be lost due to natural wear or breakage of the tip of the tusk. According to Hay (1984) and references cited therein, 17.8%–34% of tusks are broken. Therefore, the change in growth rate at 4–6 years of age could help to identify the approximate number of missing GLGs (by locating the change in growth-rate slope; Fig. 2a,b).

## 5 Conclusions

This study is the first to attempt non-destructive tusk ageing and direct measurement of age-specific growth rates in narwhal teeth using CT. The results show that (1) the growth of erupted tusks does not simply decrease with age, but peaks in adolescence; and (2) the growth rates of embedded and erupted teeth diverge at the onset of adolescence, presumably reflecting a differential tissue-level response to sexual maturation (Garde et al., 2007; Graham et al., 2020; Hay, 1984; Watt et al., 2020). The study would benefit from a larger sample size, which is not available to the authors. However, the results demonstrate that CT can be used to estimate the age of precious private and museum narwhal specimens, which might lead to insights that benefit the sustainable harvest of this endemic whale. Without additional blind tests and when all subjective and objective methods do not give exactly the same age (Reiter et al., 2025b), it remains unclear how accurate these estimates are. We suggest that the accuracy of the CT estimates might be comparable to or better than that of visual GLG counts, because the reader is free from the main sources of error (i.e., a section is not mid-longitudinal or stain and decalcification are incorrect; (Hohn, 2018)). Over the last four decades of research, only *<*130 tusked narwhals have been aged and their tusk lengths measured (Dietz et al., 2021; Garde et al., 2012, 2007; Hay, 1984; Reiter et al., 2025b). Given that museums and private collections around the world contain a large number of specimens (Nweeia et al., 2012), CT provides a key to obtaining age data that was previously inaccessible. In addition, the data generated in this and similar follow-up studies could be used for 3D printing of narwhal teeth or skulls, which would be valuable for educational and outreach purposes.

## Acknowledgments

We thank M. Eda and T. Tsubota for the opportunity to scan the specimens, and M. Otsuki, S. Sugiyama, R. Kusaka, T. Mori, and T. Oshima for supporting their preparation and transport to Japan. Specimens from other narwhals were kindly provided by M. Ogawa (NW2313, NW24, NW2403). A. Stallard improved the English in the manuscript. We also thank J. Teilmann, C. Watt, and R. Dietz for help in finding literature, and H. Birkedal and other organizers for the invitation to present at the international symposium “Narwhal tusks: a tale with a twist” (8-10 June 2026, Copenhagen, Denmark), which partly motivated this study.

## Author contribution

EAP: Conceptualization, Data curation, Formal analysis, Investigation, Methodology, Resources, Supervision, Visualization, Writing - original draft. GS: Methodology, Resources, Writing - review & editing. SMP: Methodology, Validation, Writing - original draft.

## Data and Code Availability

The generated data are available in the Zenodo repository (Podolskiy et al., 2026). We did not generate any new code in this study. Data processing and visualizations were made with freely available Weasis v. 4.6.6, 3D Slicer v. 5.10.0, and ImageJ v. 1.54g, and commercially available Matlab R2022b.

